# Early-life language deprivation affects specific neural mechanisms of semantic representations

**DOI:** 10.1101/2022.11.06.515336

**Authors:** Xiaosha Wang, Bijun Wang, Yanchao Bi

## Abstract

One signature of the human brain is its ability to derive knowledge from language inputs, in addition to nonlinguistic sensory channels such as vision and touch. How does human language experience specifically modulate the way in which semantic knowledge is stored in the human brain? We investigated this question using a unique early-life language-deprivation human model: early deaf adults who are born to hearing parents and thus had delayed acquisition of any natural human language (speech or sign), with early deaf adults who acquired sign language from birth as nonlinguistic sensory experience controls. Neural responses in a meaning judgment task with 90 written words that were familiar to both groups were measured using fMRI. The early language-deprived group, compared with the deaf control group, showed reduced semantic sensitivity, in both multivariate pattern (semantic structure encoding) and univariate (abstractness effect) analyses, in the left dorsal anterior temporal lobe (dATL). These results provide positive, causal evidence that the neural semantic representation in dATL is specifically supported by language, as a unique mechanism of representing (abstract) semantic space, beyond the sensory-derived semantic representations distributed in the other cortical regions.

## Introduction

Humans are believed to be the only species in the animal kingdom where knowledge learning can be achieved symbolically, mostly through language (“roses are red”), in addition to sensory channels (visually perceiving roses in color red) (Gelman and Roberts, 2017; Perszyk and Waxman, 2018). A common view shared by the modern neurocognitive theories of semantics is that semantic knowledge, even those acquired through language, is ultimately grounded in (nonlinguistic) sensory/motor experiences, encoded in the distributed brain areas encompassing high-level sensorimotor areas, and potential hub regions that bind such sensory-derived representations (Barsalou, 2016; Binder et al., 2009; Fernandino et al., 2022; Lambon Ralph et al., 2017; Martin, 2016). Are there neural structures that specialize to represent knowledge derived from language experience, and not (nonlinguistic) sensory experience?

Neural representations of fully nonsensory, language-derived knowledge have only recently been inferred based on studies with sensory-deprived individuals (Bottini et al., 2020; Striem-Amit et al., 2018; Wang et al., 2020, see (Bi, 2021) for a review). Congenitally blind individuals, who cannot acquire visual-specific knowledge (e.g., color) through sensory experiences, can nevertheless acquire semantic structures about such knowledge behaviorally similar to those in the sighted, presumably derived from language. Such nonsensory semantic structures in the blind (and also sighted) are represented in the dorsal anterior temporal lobe (dATL), with sighted individuals additionally representing the highly similar knowledge structure (presumably derived from sensory experience) in the visual cortex (Wang et al., 2020). This dATL cluster is distinct from the central “amodal” semantic hub proposed to bind together multiple sensory attributes, which lies in the more ventral-medial territory of the ATL (Lambon Ralph et al., 2017; Patterson et al., 2007; see discussions in Striem-Amit et al., 2018). The left dATL is also more strongly activated by abstract than concrete words in typically developed individuals (Binder et al., 2009; Bucur and Papagno, 2021; Wang et al., 2010, 2019), corroborating the role of this area in representing knowledge derived from language (see discussions in Bi, 2021). These lines of evidence, while highly suggestive, are based on manipulating sensory experience by examining individuals deprived of a sensory modality or by contrasting concepts with rich sensory experiences versus those without, rather than the positive manipulation of language experience. Thus the positive evidence for the necessity of language experiences for the neural semantic representation here is still lacking.

Here, we take advantage of a special early-life language-deprivation human model: individuals who were born profoundly deaf in hearing families and thus had very limited natural language exposure (speech or sign) during the critical period of language acquisition in early childhood (Goldin-Meadow and Feldman, 1977; Mayberry et al., 2002). These individuals usually acquired their first language (sign language) around school age, which is much later than the age of first language (L1) acquisition in typically developed children or native deaf signers who were born in deaf families and acquired sign language from birth. Previous research has shown that these delayed deaf signers have lower proficiency in phonological, morphological, and syntactic processing of sign language (Bogliotti et al., 2020; Caselli et al., 2021; Cheng and Mayberry, 2021; Lieberman et al., 2015; Mayberry et al., 2002; Mayberry and Fischer, 1989; Newport, 1990; Tomaszewski et al., 2022) and decreased neural activation in language areas (Mayberry et al., 2018, 2011; Richardson et al., 2020; Twomey et al., 2020) even after many years of sign language usage, indicating long-lasting hypofunction of the language system as a result of missing the critical period of language acquisition. The effects of early language deprivation on semantic processing have been scarcely studied and the extant pieces of evidence reported minimal influences in semantics-related behavioral or N400 measures in adult signers (Baus et al., 2008; Davidson and Mayberry, 2015; Skotara et al., 2012). Given that semantic processing is supported by a multifaceted cognitive system and a complex neural network entailing distributed semantic regions (Bi, 2021; Binder and Desai, 2011; Lambon Ralph et al., 2017; Martin, 2016), the comparable behavioral performances do not indicate comparable neural structures (e.g., color knowledge in the blind and sighted people (Wang et al., 2020)). We compared native and delayed deaf signers in terms of their fMRI BOLD responses to meaning judgment to examine their neural substrates of semantic processing of words that are familiar to them. That is, by a rare opportunity of manipulation of early language experience offered by nature, we empirically examined the specific role of language experience in the semantic neural representation.

## Results

### Participants’ background information and task fMRI design

We recruited two adult groups of congenitally or early deaf participants (Table 1), including 16 with native sign language acquisition (native deaf signers) and 23 with delayed language acquisition (delayed deaf signers). Native signers were born in deaf families; delayed signers were born in hearing families and became exposed to Chinese sign language (CSL) between the ages of 4 and 10 (mean ± standard deviation (SD): 6.91 ± 1.62 years old). The family signing environment was confirmed by the subjective ratings of parental CSL proficiency: Native signers rated their parents to have much more proficient CSL skills than delayed signers did on a 7-point scale (Welch’s *t*_*31*.*7*_ = 13.54, *p* = 1.10 × 10^−14^; each participant provided CSL ratings for both parents and the maximum score was used for group comparison). All deaf participants received formal education in special education programs since elementary school. The two deaf groups were matched on demographic variables (gender, age, years of education, *p*s > .15) and subjective social status (Adler et al., 2000) during childhood (*p* = .54). While the two groups significantly differed in the education levels of their parents (Table 1; *p*s < .053), the direction was in favor of delayed signers as their hearing parents received more formal education than the deaf parents of native signers. In terms of language skills, the two deaf groups were matched on self-reported proficiency of CSL comprehension, production, and lipreading skills (*p*s > .34), and on the reading disability risk measured by the adult reading history questionnaire (Lefly and Pennington, 2000) (*p* = .22). Written word processing was further evaluated in two reaction-time tasks (i.e., visual lexical decision and word-triplet semantic judgment) and no significant group differences were found (Supplementary file 1). In summary, the native and delayed deaf groups were carefully matched on a wide range of demographic and later language performances (sign, lipreading, and written). Thus the early language deprivation is a strong candidate to account for the neural group differences reported below.

**Table 1.** Background demographic and language information of native and delayed signers.

|  | native<br>(n = 16) | delayed<br>(n = 23) | Welch's<br>t | p-value | Hedges'<br>g |
| --- | --- | --- | --- | --- | --- |
| AoA of CSL (years old) | 0 ± 0 | 6.91 ± 1.62 |  |  |  |
| Parental CSL proficiency (1-7 scale) <sup>a</sup> ** | 6.81 ± 0.54 | 2.74 ± 1.29 | 13.54 | <.001 | 4.04 |
| Gender | 11M, 5F | 12M, 11F |  |  |  |
| Age (years) | 28.50 ± 7.13 | 27.09 ± 5.87 | 0.65 | .52 | 0.21 |
| Education (years) | 14.13 ± 2.31 | 15.09 ± 1.41 | -1.49 | .15 | -0.49 |
| CSL comprehension (1-7 scale) <sup>a</sup> | 6.06 ± 0.93 | 6.17 ± 0.89 | -0.38 | .71 | -0.12 |
| CSL production (1-7 scale) <sup>a</sup> | 5.75 ± 1.00 | 5.87 ± 1.10 | -0.35 | .73 | -0.11 |
| Lipreading of acquaintances (1-7 scale) <sup>a</sup> | 2.81 ± 1.11 | 3.17 ± 1.19 | -0.97 | .34 | -0.31 |
| Lipreading of strangers (1-7 scale) <sup>a</sup> | 1.88 ± 0.81 | 1.87 ± 0.97 | 0.02 | .99 | 0.01 |
| Adult Reading History Questionnaire (0-4 scale) <sup>b</sup> | 2.59 ± 0.68 | 2.84 ± 0.52 | -1.25 | .22 | -0.41 |
| McArthur Scale of Subjective Social Status, childhood (1-10 scale) <sup>c</sup> | 3.88 ± 2.58 | 4.35 ± 1.97 | -0.62 | .54 | -0.20 |
| Father's education (years) | 7.19 ± 3.04 | 10.39 ± 2.84 | -2.02 | .053 | -0.65 |
| Mother's education (years)** | 6.56 ± 3.14 | 8.52 ± 2.73 | -3.33 | .002 | -1.07 |

Notes:
a: higher scores indicate higher proficiency.
b: higher scores indicate higher risk of reading disability; from (Lefly and Pennington, 2000).
c: higher scores indicate higher social status; from (Adler et al., 2000).
\*\*: $p < .01$ , significant differences between 2 deaf groups.

The stimuli of task fMRI were 90 written words that were highly familiar to both groups, including 40 concrete/object words varying in sensory and motor properties (animals, face/body parts, artefacts) and 50 abstract/nonobject words varying in social and emotional contents. These words could be grouped into 10 categories based on the group-averaged semantic multi-arrangement performances in an independent group of 32 hearing participants (see Methods, Figure 1a, and Figure 1-figure supplement 1). Both deaf groups judged these words to be highly familiar (7-point familiarity ratings from 8 native signers: 6.72 ± 0.17; 13 delayed signers: 6.86 ± 0.15; 7 being most familiar) and yielded similar word-relational structures in a 1-hour semantic distance judgment task, in which each participant (16 native and 21 delayed signers) produced a 90 × 90 semantic distance matrix (RDM). The group-averaged semantic RDMs of the two deaf groups were strongly correlated (Spearman rho = 0.71, n = 4005; Figure 1a). At the individual level, correlations with the benchmark (the group-averaged RDM of hearing participants) did not significantly differ between the two deaf groups (Welch’s *t*_*31*.*30*_ = -0.37, *p* = .71).

**Figure 1.**
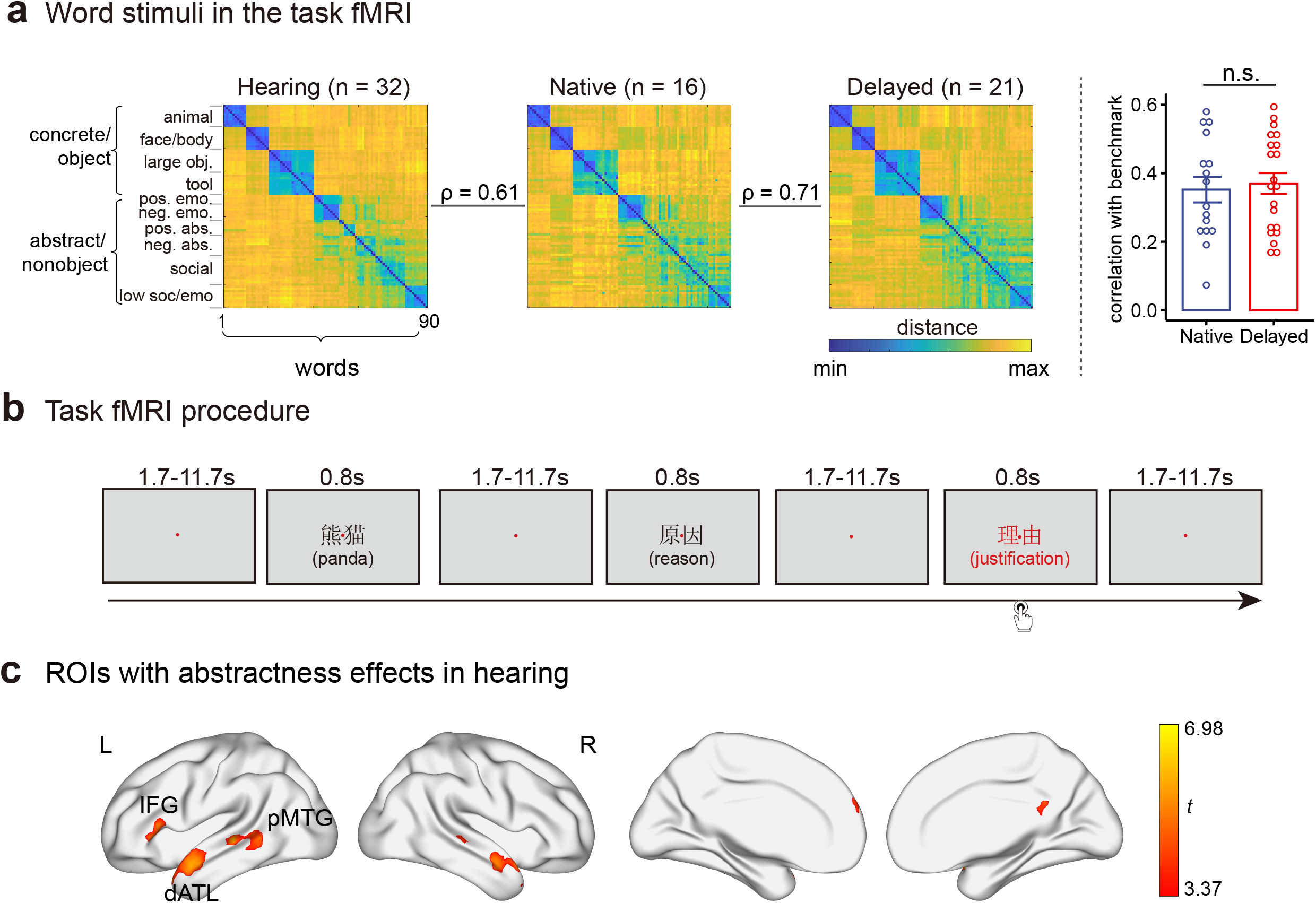
The word stimuli, task fMRI procedure, and ROIs in this study. **a**. Word stimuli in the task fMRI. Ninety words were used, including 40 concrete/object words and 50 abstract/nonobject words, which were grouped into fine-grained categories based on k-means clustering of the group-averaged semantic space of 32 hearing participants. The left panel shows the group-averaged semantic representational matrices (RDM) in hearing participants, native signers, and delayed signers. The right panel shows the Spearman’s rho between each deaf participant’s semantic RDM and the group-averaged semantic RDM in hearing participants. **b**. Task fMRI procedure. During scanning, participants were asked to think about each of 90 target word meanings (displayed in black, e.g., “panda”, “reason”) and to determine whether occasional words displayed in red (catch trials, e.g., “justification”) were semantically related to the previous word. There were 90 target word trials (each word appeared once) and 14 catch trials in each run. **c**. Semantic ROIs were functionally identified by contrasting abstract/nonobject words with concrete/object words in 33 hearing participants at the threshold of voxel-level *p* < .001, cluster-level FWE-corrected *p* < .05. dATL, dorsal anterior temporal lobe; IFG, inferior frontal gyrus; pMTG, posterior middle temporal gyrus. L, left hemisphere; R, right hemisphere. The following source data and figure supplement are available for figure 1: **Source data 1:** Spearman’s rho between each deaf participant’s semantic RDM and the group-averaged semantic RDM in hearing participants. **Figure supplement 1**. Ninety words in the fMRI task, grouped into ten semantic categories. **Figure supplement 2**. Details of semantic ROIs, including cluster location, extent, peak t values, and MNI coordinates. **Figure supplement 3**. RSA and univariate results in other ROIs (except for the left dATL, IFG, and pMTG) in two deaf groups.

In the MRI scanner, participants were asked to think about the meaning of each of the 90 words (condition-rich fMRI design (Kriegeskorte et al., 2008a)) and to decide whether a word in red (catch trials) was semantically related to its previous word (i.e., oddball one-back semantic judgment, Figure 1b). We first examined the two-deaf-group differences in brain regions preferring abstract/nonobject words to concrete/object words, which have been proposed to relate to verbal semantic processing (Binder et al., 2009; Wang et al., 2010). These regions were functionally localized by contrasting abstract/nonobject words (e.g., *reason*) to concrete/object words (e.g., *panda*) in an independent group of 33 hearing participants (voxel-level *p* < .001, cluster-level FWE-corrected *p* < .05). This contrast in the hearing group resulted in regions in the frontal and temporal cortices that were well aligned with the literature (Figure 1c and Figure 1-figure supplement 2). In particular, the left dATL, the left inferior frontal gyrus (IFG), and the left posterior middle temporal region (pMTG) were most consistently reported in the meta-analyses (Binder et al., 2009; Bucur and Papagno, 2021; Wang et al., 2010) and were taken as our primary regions of interest (ROIs). Results for the other clusters are shown in Figure 1-figure supplement 3 (no significant group differences). We then carried out whole-brain analyses to explore the two-deaf-group differences beyond these semantic ROIs. We focused on two neural semantic effects: 1) multivoxel pattern representation of word meaning (i.e., semantically related words have similar neural patterns); and 2) the univariate semantic abstractness effects. It should be noted that the neural semantic abstractness effect does not equate with language-derived semantic knowledge, as it might arise from some nonverbal cognitive processes that are more engaged in abstract word processing (Binder et al., 2016). The reasoning here is that if a brain region’s abstractness effect is affected by early language deprivation, this would constitute further evidence that the abstractness computed by this brain region is indeed associated with language processes.

### Semantic structure representation: dATL alteration in delayed signers

We first examined whether early-life language deprivation affects neural representations of the semantic space using representational similarity analysis (RSA, Figure 2a) (Kriegeskorte et al., 2008a, 2008b). We estimated the multivoxel activation pattern for each target word and, in a given ROI, calculated the correlation distance (1-Pearson correlation) of the multivoxel activation patterns between each word pair to build a 90 × 90 neural representational dissimilarity matrix (RDM). As a sanity check, we first carried out whole-brain searchlight RSA of visual similarity of written words, by computing Spearman’s rank correlation between the pixelwise dissimilarity of the 90 words and neural RDMs, and observed significant encoding of written word visual similarity in the early visual cortex in both native and delayed signers at the threshold of voxel-level *p* < .001, cluster-level FWE-corrected *p* < .05 (Supplementary file 2). To quantify the semantic information encoded in the neural RDMs, we computed Spearman’s partial correlation between the neural RDMs and the 10-category benchmark semantic RDM (i.e., those of the hearing group), while controlling for the stimulus low-level visual and phonological RDMs.

**Figure 2.**
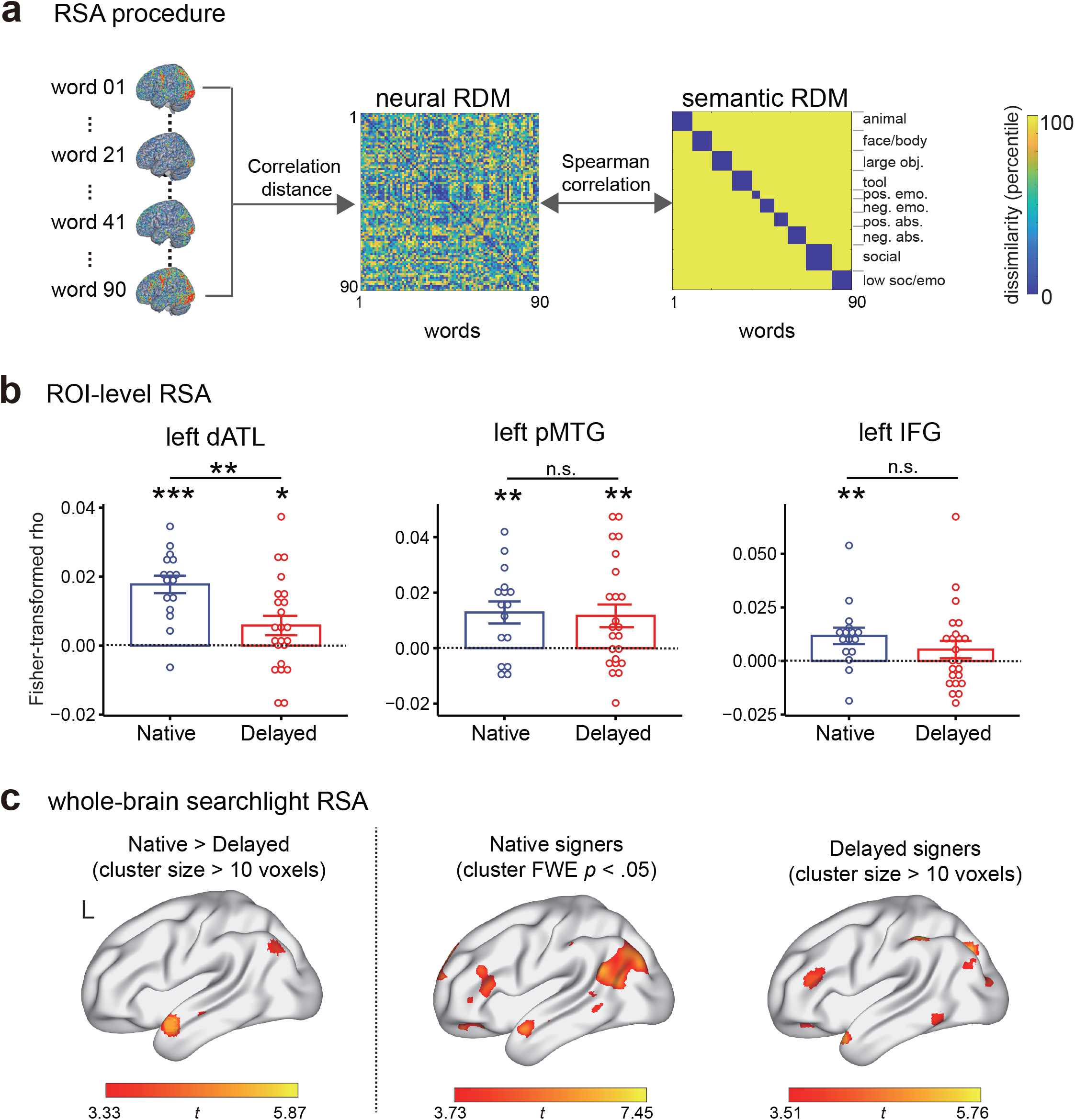
Effects of early language deprivation on neural representations of semantic knowledge. **a**. RSA procedure. For each participant, a neural RDM was computed as the correlation distance of multivoxel activity patterns (in a given ROI) for each pair of words and then correlated with the hearing-group-level semantic category RDM to quantify semantic information encoded in the neural RDM. **b**. ROI-level RSA results. Error bars indicate 1 s.e.m. n.s., not significant, *p* > .05; *, *p* < .05; **, *p* < .01; ***, *p* < .001; **c**. Whole-brain searchlight RSA results. The statistical maps were thresholded at voxel-level *p* < .001, cluster FWE-corrected *p* < .05, for native signers and at voxel-level *p* < .001, cluster size > 10 voxels, for delayed signers and group comparisons. Brain results of group comparisons and delayed signers were visualized using the “Maximum Voxel” mapping algorithm in BrainNet Viewer to illustrate small clusters. Clusters in the left hemisphere are shown here. n = 16 in the native group; n = 23 in the delayed group. The following source data and figure supplement are available for figure 2: **Source data 1:** ROI-level RSA results in two deaf groups. **Figure supplement 1**. Cluster details of whole-brain searchlight RSA results.

ROI-level RSA was carried out in the left dATL, pMTG, and IFG. As shown in Figure 2b, the three ROIs significantly encoded the semantic space in native signers (*t*_15_ > 3.04, one-tailed *p*s < .004, Cohen’s d > 0.76). In delayed signers, the left dATL and pMTG significantly encoded semantic space (*t*_22_ > 2.07, one-tailed *p*s < .025, Cohen’s d > 0.43) and the left IFG showed a similar, nonsignificant, trend (*t*_22_ = 1.31, one-tailed *p* = 0.10, Cohen’s d = 0.27). Critically, group differences were observed in the left dATL (Welch’s *t*_36.7_ = 3.15, two-tailed *p* = .003, Hedges’ g = 0.98), not in the left pMTG or IFG (*p*s > .26). For the left dATL, we further computed a partial correlation between neural RDMs and the semantic RDM, controlling for the object-vs-nonobject category structure. The reduction in the delayed group compared with native signers still held (Welch’s *t*_30.4_ = 2.91, two-tailed *p* = .007, Hedges’ g = 0.93), indicating that language-deprivation-induced semantic-encoding downregulation was beyond the object-vs-nonobject binary distinction in the left dATL.

We then carried out whole-brain searchlight RSA (Kriegeskorte et al., 2006) to explore group differences in semantic encoding beyond the semantic ROIs defined above (Figure 2c and Figure 2-figure supplement 1). At the threshold of voxel-level *p* < .001, cluster-level FWE-corrected *p* < .05, in native signers, the semantic encoding was observed in the left dATL, IFG, pMTG, along with adjacent parietal and occipital areas, dorsomedial prefrontal cortex, and frontal orbital cortex. In delayed signers, no brain regions survived the whole-brain correction; at a more lenient threshold of voxel *p* < .001, cluster size > 10 voxels, semantic encoding could be observed in the left dATL, IFG, and clusters scattered in the left inferior parietal and occipital regions. Importantly, the whole-brain contrast between the two deaf groups peaked at the left dATL at the threshold of voxel *p* < .001, which was also the largest cluster (Brodmann area 21/38, peak MNI coordinates: -60, 6, -20, peak t = 5.87, 162 voxels), converging well with the ROI results in that native signers showed stronger semantic encoding than delayed signers in the left dATL ROI. While this cluster was not large enough to survive the whole-brain correction (cluster FWE-corrected *p* = .22), the top 10 voxels survived the voxel-level correction of FWE-corrected *p* < .05. No areas were found to show significantly increased semantic information in delayed signers compared with native signers at the conventional threshold.

### Univariate semantic abstractness effects: dATL alteration in delayed signers

We then examined how early language deprivation might alter the neural semantic abstractness effects by comparing regional activation strength to abstract/nonobject and concrete/object words in two deaf groups. At the ROI level, all three ROIs (the left dATL, IFG, and pMTG) exhibited significant abstractness preference in both deaf groups (Figure 3a; native group: paired *t*s > 3.72, df = 15, one-tailed *p*s < .001, Cohen’s *d*s > 0.93; delayed group: paired *t*s > 2.13, df = 22, one-tailed *p*s < .02, Cohen’s *d*s > 0.44), further indicating the robustness of semantic abstractness preference in these ROIs. We then carried out two-way ANOVA to examine group effects in each of these ROIs, with word type (concrete/object, abstract/nonobject) as a within-subject factor and group (native, delayed) as a between-subject factor. The group × word-type interaction reached statistical significance in only the left dATL (*F*_*(1,37)*_ = 4.91, *p* = .033), not in the left pMTG or IFG (*ps* > .30). In the left dATL native signers exhibited greater semantic abstractness effects than delayed signers. The main effects of group were not significant in any of the three ROIs (*p*s > .057).

**Figure 3.**
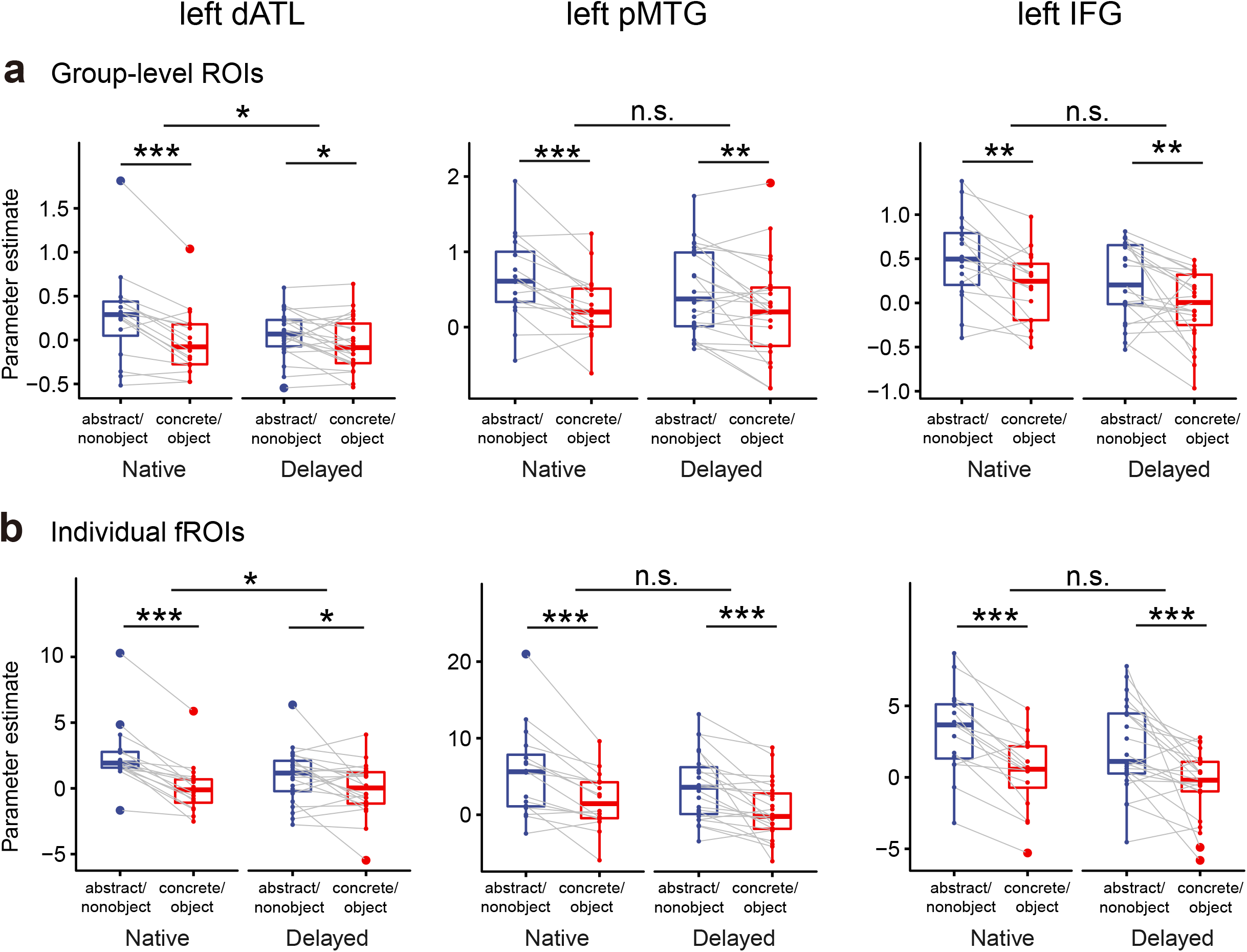
Effects of early language deprivation on the univariate semantic abstractness effects in group-level-defined semantic ROIs (**a**) and individual fROIs (**b**). Boxplots show beta values to abstract/nonobject words and concrete/object words in the three ROIs (the left dATL, pMTG, and IFG). n.s., not significant, *p* > .05; *, *p* < .05; **, *p* < .01; ***, *p* < .001. n = 16 in the native group; n = 23 in the delayed group. The following source data and figure supplement are available for figure 3: **Source data 1:** Raw beta values to abstract/nonobject words and concrete/object words in group-level-defined semantic ROIs and individual fROIs in two deaf groups. **Figure supplement 1**. Whole-brain univariate results of semantic abstractness in native and delayed signers.

Considering inter-subject variations in activation locations, we further carried out the individualized functional region-of-interest (fROI) analysis for validation (Cohen et al., 2019; Murty et al., 2020) (see Methods). Using each of the ROIs defined in the hearing group as anatomical constraints, we identified the top 50 selective voxels in each deaf participant using one-half of the fMRI data and computed regional activation strength to abstract and concrete words in these voxels in the held-out data (i.e., via an odd-even run cross-validation process). As shown in Figure 3b, the group × word-type interaction was again observed in the left dATL (*F*_*(1,37)*_ = 6.20, *p* = .017), not in pMTG or IFG (*p*s > .58); main effects of group were not found (*p*s > .18).

We further carried out a whole-brain analysis of the semantic abstractness effects in the two deaf groups (Figure 3-figure supplement 1). At the threshold of voxel-level *p* < .001, cluster-level FWE-corrected *p* < .05, semantic abstractness reached significance in the left dATL in native signers (Brodmann area 21/38/22, peak MNI coordinates: -58, 4, -8, peak t = 7.27, 209 voxels). No regions survived the whole-brain threshold in delayed signers; at a lenient threshold of voxel *p* < .001, cluster size > 10 voxels, abstractness can be observed in the left pMTG and dATL, which are consistent with the ROI results above. For the whole-brain comparisons of the semantic abstractness effects between the two deaf groups, no areas survived the conventional threshold; small scattered clusters (cluster sizes < 14 voxels) were observed at a lenient threshold of voxel *p* < .001.

## Discussion

By comparing fMRI BOLD responses to the word semantic structures between native and delayed early deaf signers, we aimed to examine how the critical-period, early-life language experience affects the semantic neural representations. Our results demonstrated that semantic information in the left dATL was significantly reduced in delayed signers compared with native signers in both multivariate (representation of rich semantic space) and univariate (preference to abstract/nonobject words) analyses. That is, early-life language acquisition—critical to language system neurodevelopment—is necessary for the left dATL to exhibit the typical semantic organization in adulthood. Such evidence provides the first positive evidence for the effect of language experience in driving the semantic functional development of a particular brain region.

First, what kind of variables best account for the current group contrast results? The two deaf groups were matched on sensory-experiences (both similarly profound deaf), a wide range of nonlinguistic environmental variables including the socioeconomic status (except for parents’ education backgrounds, which were better for the delayed signer group), the length and level of formal education, through which largely comparable sign language and word reading skills were achieved. The salient difference was the early-life language experience (before 6.9 years old -- the mean age of acquisition in our delayed signer group). Notably, deaf children with early language deprivation often spontaneously develop homesign systems, which share some aspects with natural language (Goldin-Meadow, 2003). For instance, they could use homesigns to produce generic utterances, more so for animals than for artefacts (Goldin-Meadow et al., 2005). Critically, our positive results of the robust group differences in dATL suggest that early homesign and later formal education for sign and written language experiences are insufficient for typical dATL neurodevelopment; the full-fledged language experience during the critical period (before school age) plays a necessary role in this process. A further variable to consider is that early language deprivation may lead to alterations beyond the language system, affecting non-language cognitive domains such as working memory or theory of mind in children (Marshall et al., 2015; Richardson et al., 2020). While we cannot rule out the effects of these potential intermediate cognitive processes, the positive evidence of these variables in dATL functionality is lacking. The most direct manipulation and parsimonious account for the dATL effects is language related (see below).

Importantly, the specific effects of language experience in shaping the semantic representation in dATL observed here are corroborated by several previous lines of indirect evidence for the relevance of language in its functionality. It shows a functional preference for abstract words, which are less salient in sensory attributes than concrete words, and are assumed to entail more language processes such as context diversity (e.g., Binder et al., 2009; Bucur and Papagno, 2021; Wang et al., 2010). In congenitally blind individuals, it encodes visual knowledge such as color and is negatively modulated by perceptibility (Bottini et al., 2020; Striem-Amit et al., 2018; Wang et al., 2020). It was further shown recently that left ATL encodes specific higher-order statistical relations (i.e., graph) among words computed from word co-occurrences patterns within the language system (Fu et al., 2022). It has also been shown to be sensitive to semantic composition, such as in two-word phrases or sentences (Pallier et al., 2011; Pylkkänen, 2019). This region is functionally and structurally connected with other language-sensitive regions (Fan et al., 2014; Friederici et al., 2017; Lambon Ralph et al., 2017; Pascual et al., 2015; Wang et al., 2019). Finally, neurodevelopmental evidence also highlighted the importance of dATL in early language and semantic development. A recent study (Yu et al., 2021) showed that the resting-state functional connectivity patterns of the middle portion of the left temporal pole in infancy and toddlerhood significantly predict performances in tasks entailing semantic processes (i.e., the Picture Vocabulary and Oral Comprehension tests), not in tasks of phonological skills or rapid automatized naming, at about 6.5 years old; this prospective association is modulated by early language skills, as controlling for early language skills significantly reduces the association. This compelling neurodevelopmental pattern indicates the dATL-related network as a neural scaffold for subsequent accumulation of language-related semantic knowledge. These lines of findings are often correlational evidence for the language effect on their own. Together with the positive evidence of early language experience in developing typical semantic representation in dATL reported here, the parsimonious proposal accounting for them together would be that dATL represents knowledge derived from language (e.g., from higher-order word relations), and that nonsensory, abstract meanings are more strongly supported by such language information. With critical period language deprivation, the semantic structure representation and the abstract preference here are reduced.

Interestingly, the semantic effects did not significantly differ between the two deaf groups in the left IFG and pMTG, two other brain regions widely implicated in language and semantic processing, also with an abstractness preference (Binder et al., 2009; Friederici et al., 2017; Lambon Ralph et al., 2017; Wei et al., 2012; Xu et al., 2017). In the early-language deprivation model, previous studies found that delayed signers show reduced activity in posterior temporal regions and/or IFG in multiple sign language sentence judgment tasks (Mayberry et al., 2011; Twomey et al., 2020) and increased IFG activity in phonological judgment (MacSweeney et al., 2008). Reduced fractional anisotropy (FA) values in the left dorsal arcuate fasciculus structurally connecting the two regions have been reported in three cases with post-childhood first-language acquisition (Cheng et al., 2019). These results indicate that the development of the dorsal language pathway, particularly those substrates supporting the syntactic and/or phonological processes, is indeed shaped by early language experience. Our results show that the semantic processes in these regions were not affected by the changes in nonsemantic language functions (e.g., phonology). While the detailed functional relations among these regions and dATL are unclear, the relatively salient role of ATL in language-supported semantics is consistent with the dynamic causal modeling results that identified the ATL (not IFG, MTG, or inferior parietal areas) as the early hub (within 250 ms) for abstractness effects, followed by inferior parietal areas as a late hub, across three electro/magnetoencephalography tasks (Farahibozorg et al., 2022). Therefore, while other brain regions may also contribute to language-supported semantics, a detailed neuroanatomical framework is needed to accommodate why language-related semantic effects peak in the dATL.

What are the implications of the current findings for neurocognitive models of semantics and semantic behaviors? The relationship between the current neural results and semantic behaviors is intriguing. On the one hand, these findings about how language experience is critical for specific semantic neural structures (dATL) are in line with the rich evidence and discussion in the behavioral literature about the role of language in semantic development. For instance, developmental behavioral evidence shows that young human infants are sensitive to language labels and statistical patterns in forming semantic categories and other types of semantic relations (Perszyk and Waxman, 2018; Unger and Fisher, 2021; Spelke, 2017; see similar discussions in Gelman and Roberts, 2017; Lupyan et al., 2020). On the other hand, consistent with previous behavioral studies of semantic processing in delayed adult signers (Baus et al., 2008; Davidson and Mayberry, 2015; Skotara et al., 2012), in our study native and delayed signers exhibited similar semantic space for words that they are familiar with (Figure 1a) and did not significantly differ in two additional reaction time tasks (Supplementary file 1). That is, we did not see visible semantic behavioral differences despite the reduced dATL semantic representation. This seeming conflict is likely related to the multifaceted nature of the cognitive and neural basis of semantics more broadly. Semantic processing entails representations that can be derived from multiple types of experiences, supported by highly distributed neural systems (e.g., Binder et al., 2009). In fact, many neurocognitive semantic theories mainly focus on the representing, binding, and controlling of those knowledge derived from nonlinguistic multisensory experiences, and are less committed about the kind of language-derived neural semantic representations (Binder, 2016; Lambon Ralph et al., 2017; Martin, 2016; but see Mahon and Caramazza, 2008). The two deaf groups are not assumed to differ in these nonlinguistic aspects of semantic processing. The delayed signers may thus rely on these systems to achieve comparable performances in the semantic tasks being tested. Our current findings that positively show the specific effect of language experience on dATL and not on other regions are thus best accommodated by a recently developed dual-neural-coding framework for semantic knowledge (Bi, 2021), which explicitly proposes that the human brain has developed specialized brain regions (the dATL) for language-derived semantic representations, in addition to the sensory-derived knowledge representations. Now with stronger evidence for the language-driven representation, future studies are warranted to examine the exact computational principles in the left dATL (e.g., Fu et al., 2022) and semantic behaviors more targeted at the functionality of this area than the general semantic tasks currently employed.

To conclude, by manipulation of early language experience by nature we were able to positively identify a human brain region – dorsal anterior temporal lobe -- specifically supporting word meanings derived from language experience, in contrast to those grounding nonlinguistic, sensory experiences. These findings highlight the unique role of language in forming specific neural semantic representations in the human brain, and raise new questions about the ontogenetic mechanisms of this intriguing neural structure.

## Materials and Methods

### Participants

Thirty-nine congenitally or early deaf adults and 33 hearing college students (15 males, mean age 21.97 ± 2.58 years, range: 18-28 years, all native Mandarin Chinese speakers) were recruited for the fMRI study. All except for 1 delayed deaf signer and 1 hearing participant participated in the behavioral semantic distance judgment task. The behavioral data of one delayed signer were excluded from data analysis due to technical errors. The behavioral and neural data of 21 out of the 33 hearing college students overlapped with a previous study (Wang and Bi, 2021) that examined the intersubject variability of semantics in typically developed populations.

Deaf participants included two groups (Table 1). Native signers (*n* = 16; 11 males) had deaf parents and were exposed to CSL soon after birth. Delayed (nonnative) signers (*n* = 23; 12 males) were born in hearing families and learned CSL after they were enrolled in special education schools (range: 4-10 years old). All deaf participants completed a background questionnaire, in which they reported their hearing loss conditions, history of language acquisition, and educational background. They were also asked to rate sign language abilities of their family members and to rate their own sign and lipreading abilities using a 7-point scale: 1 = none at all, 7 = very proficient. All deaf participants then finished the Chinese version of the Adult Reading History Questionnaire (a self-report screening tool for the risk of reading disability in adults (Lefly and Pennington, 2000)), the MacArthur Scale of subjective social status (Adler et al., 2000) during childhood, and reported the educational years of their parents. For hearing loss conditions, all deaf participants reported being severely or profoundly deaf from birth, except for 1 native signer and 3 delayed signers who reported becoming deaf before the age of 2 due to medication side effects. Hearing thresholds in decibels (dB) were available in 23 deaf participants (native: 11/16; delayed: 12/23) and confirmed severe to profound hearing loss (range: 85-120 dB). One native signer and 4 delayed signers were using hearing aids at the time of testing; others either had never used hearing aids (6 native signers and 5 delayed signers) or had used hearing aids for some time (9 native signers and 14 delayed signers; years of use: 0.5-20 years). Speech comprehension was reported to be very poor, even with the use of hearing aids.

All participants had normal or corrected-to-normal vision. All participants were right-handed, except for three native signers (1 was ambidextrous and 2 were left-handed) (measured by the Edinburgh inventory (Oldfield, 1971)). All participants gave written informed consent and received monetary compensation for participation. The study was approved by the Human Subject Review Committee at Peking University (2017-09-01), China, in accordance with the Declaration of Helsinki.

### Word stimuli in task fMRI

The stimuli for fMRI scanning were 90 written words, including 40 concrete/object words and 50 abstract/nonobject words without explicit external referents. Concrete/object words varied in sensory and motor attributes and included 10 animals (e.g., *panda*), 10 face or body parts (e.g., *shoulder*), 10 tools (e.g., *hammer*), and 10 common large household objects (e.g., *microwave*). Abstract/nonobject words varied in social and emotional associations (e.g., *cause, violence*). A 7-point familiarity rating, with 7 being the most familiar, was collected from 21 out of 39 deaf signers and an independent group of 26 hearing college students. All words were highly familiar to deaf participants (8 native signers: 6.72 ± 0.17; 13 delayed signers: 6.86 ± 0.15). All words were disyllabic except for five object words (“cat” and “bed” are monosyllabic and “giraffe”, “microwave”, and “washing machine” are trisyllabic in Chinese). All words were primarily used as nouns except for 10 words denoting emotional states (nine were primarily used as adjectives and one as a verb). The concrete and abstract words were matched on the number of strokes (a measure of visual complexity for Chinese words; 17.23 ± 5.80 vs. 16.14 ± 3.98; two-samples *t*_*88*_ = 1.05, *p* = .30). While concrete/object words were less frequent in a Mandarin Chinese corpus (Sun et al., 1997) than abstract/nonobject words (log word frequency: 1.05 ± 0.73 vs. 1.62 ± 0.68; *t*_*88*_ = -3.83, *p* < .001), the former were rated as more subjectively familiar than the latter in either hearing (6.79 ± 0.16 vs. 6.22 ± 0.26; *t*_*88*_ = 12.05, *p* < .001) or deaf (6.85 ± 0.07 vs. 6.77 ± 0.13; *t*_*88*_ = 3.41, *p* < .001) participants.

### Behavioral semantic distance judgment task

To examine participants’ understanding of the 90 words used in the task fMRI, we asked them to judge semantic distance among these words by arranging them spatially close together or far apart in a circular arena on a computer screen via mouse drag-and-drop operations (Kriegeskorte and Mur, 2012). The task lasted for one hour and produced a 90 × 90 semantic distance matrix (i.e., a behavioral representational dissimilarity matrix, RDM) for each participant. Taking the hearing group-averaged semantic RDM as a benchmark, we correlated each deaf participant’s semantic RDM with it using Spearman’s rank correlation, Fisher-z-transformed the correlation coefficients, and compared the two deaf groups to assess the similarity between their semantic structures.

Considering the multidimensional and flexible nature of semantic distance among various concrete and abstract words (Binder et al., 2016; Conca et al., 2021), we focused on the categorical structure of 90 words that captured the large-scale semantic relationships and was likely to be shared across individuals. The categorical RDM was obtained by performing a k-means clustering analysis on the group-averaged 90 × 90 semantic RDM of hearing participants (the factoextra package, http://www.sthda.com/english/rpkgs/factoextra, in the R programming environment; Version 4.0.0; R Core Team, 2020). The optimal number of clusters was determined based on gap statistic (Tibshirani et al., 2001), which revealed 10 semantic categories (Figure 1a and Figure 1-figure supplement 1).

### Task fMRI procedure

In the fMRI scanner, participants were instructed to look at each of the 90 target words, think about their meanings, and perform an oddball one-back semantic judgment task. In this oddball task, participants were asked to judge whether occasional words in red (catch trials) were semantically related to the previous word by pressing the corresponding buttons with the right index or middle finger.

Each participant performed 10 runs (360 s per run) of task fMRI scanning, except for one native signer who finished eight runs and withdrew due to discomfort. Each run consisted of 90 2.5 s-long target word trials and 14 2.5 s-long catch trials. Each trial started with a 0.8-s word stimulus (black color, Song font, and 2.6 visual degrees in height) at the center of a gray background, followed by 1.7-s fixation. Thirty 2.5 s-long null trials (fixation only) were randomly inserted among target and catch trials, with the interval between two words ranging from 1.7 to 11.7 s. Each target word appeared once in each run, and the order was pseudorandomized for each run in each participant. Each run began with a 12-s fixation period and ended with a 13-s rest period during which participants received a written cue that the current run was about to end. Stimulus presentation was controlled by E-prime 2. The 140 catch trials were created by first pairing each of the 90 target words with a probe word and then pseudo-randomly selecting 50 out of 90 target words (21 concrete/object words and 29 abstract/nonobject words) to pair with another 50 probe words. Participants performed this task attentively, with overall miss rates lower than 22.1% (median: 1.4%).

### Image acquisition

All functional and structural MRI data were collected using a Siemens Prisma 3T Scanner with a 64-channel head-neck coil at the Center for MRI Research, Peking University. Functional data were acquired with a simultaneous multi-slice echoplanar imaging sequence supplied by Siemens (62 axial slices, repetition time (TR) = 2000 ms, echo time (TE) = 30 ms, multi-band factor = 2, flip angle (FA) = 90°, field of view (FOV) = 224 mm × 224 mm, matrix size = 112 × 112, slice thickness = 2 mm, gap = 0.2 mm, and voxel size = 2 × 2 × 2.2 mm). A high-resolution 3D T1-weighted anatomical scan was acquired using the magnetization-prepared rapid acquisition gradient echo sequence (192 sagittal slices, TR = 2530 ms, TE = 2.98 ms, inversion time = 1100 ms, FA = 7°, FOV = 224 mm × 256 mm, matrix size = 224 × 256, interpolated to 448 × 512, slice thickness = 1 mm, and voxel size = 0.5 × 0.5 × 1 mm).

### Image preprocessing

Functional images were preprocessed using SPM12 (Wellcome Center for Human Neuroimaging, London, UK, http://www.fil.ion.ucl.ac.uk/spm12/). For each participant, the first 4 volumes of each functional run were discarded for signal equilibrium. The remaining images were corrected for slice timing and head motion and then spatially normalized to Montreal Neurological Institute (MNI) space via unified segmentation (resampling into 2 × 2 × 2 mm voxel size). All participants had head motion less than 1.97 mm/1.95°, except for one hearing participant showing excessive head motion in 2 runs (> 2 mm/2°); we thus analyzed the remaining 8 runs of fMRI data for this participant. We also estimated each participant’s framewise displacement (FD) from translations and rotations (Power et al., 2012); the two deaf groups exhibited comparable FD during scanning (Native: 0.12 ± 0.04; Delayed: 0.11 ± 0.04; Welch’s *t*_*29*.*08*_ = 0.77, *p* = .45). Spatial smoothing was applied with a Gaussian kernel of 6 mm full width at half maximum (FWHM) for univariate analysis and 2 mm FWHM for representational similarity analysis (RSA).

### ROI definition

A GLM was built to localize semantic abstractness in the brain, i.e., brain regions showing stronger activation to abstract/nonobject words than to concrete/object words, in hearing participants. For spatially smoothed functional images, the GLM included 3 regressors (i.e., abstract/nonobject words, concrete/object words, and catch trials) for each run, each convolved with the canonical hemodynamic response function (HRF). The GLM also included six head motion parameters and a global mean predictor of each run. The high-pass filter was set at 128 s. The contrast (i.e., abstract > concrete) was computed for each participant.

The brain regions of semantic abstractness were considered as candidates supporting knowledge derived from language. These regions were functionally localized in hearing participants. The beta-weight images of abstract > concrete in hearing participants were submitted to one-sample *t*-tests. A conventional cluster-extent-based inference threshold (voxel-level *p* < .001, cluster-level FWE-corrected *p* < .05) was adopted and we stated explicitly when other thresholds were applied. At the conventional threshold, six clusters were found (Figure 1c and Figure 1-figure supplement 2). As the largest cluster extended from the left dATL to the left IFG, we separated the dATL (808 voxels) and IFG (411 voxels) into two ROIs based on the automated anatomical labeling (AAL) atlas (Tzourio-Mazoyer et al., 2002). To have the cluster sizes comparable across ROIs, we increased voxel-level threshold to *p* < .0001 and obtained a smaller dATL cluster (488 voxels), based on which the ROI-level results were reported; the results were largely similar across different sizes of the dATL ROI.

### Representational similarity analysis in two deaf groups

#### GLM

The preprocessed functional images were analyzed using a GLM to create a *t*-statistic image for each word (Kriegeskorte et al., 2008a). For each participant, a GLM was built with the concatenated time series across all the scanning runs, including 90 regressors corresponding to each word and one regressor for catch trials, convolved with a canonical HRF. Additionally, 6 head motion parameters and a global mean predictor were included for each run. A high-pass filter cutoff was set as 128 s. The resulting *t*-maps for each word versus baseline were used to create neural RDMs.

#### ROI-level RSA

For each ROI in each participant, we extracted the activity patterns to each word from its whole-brain t-statistic images and calculated a neural 90 × 90 RDM based on the Pearson distance between activation patterns for each pair of words. Semantic information encoded in this ROI was quantified by computing Spearman’s rank correlation between the neural RDM and the 10-category benchmark semantic RDM. Partial correlations were used to control for two stimulus low-level property RDMs: (1) The low-level visual RDM was computed by Pearson’s correlation distance between the binary silhouette images of each word pair. (2) The word phonological RDM: Despite that our task did not require explicit phonological retrieval, there might be automatic phonological activation. We thus constructed the phonological RDM by calculating one minus the proportion of shared sub-syllabic units (onsets or rhyme) between each word pair. The resulting correlation coefficients between neural and semantic RDMs were Fisher-*z*-transformed and compared with zero using one-sample t-tests (one-tailed) to test whether the ROI significantly encoded the semantic structure. Group differences in semantic encoding were assessed using two-tailed Welch’s t-tests for each ROI.

#### Whole-brain searchlight RSA

To explore the potential differences between native and delayed signers in the neural semantic representations beyond the semantic ROIs identified above, we carried out whole-brain searchlight RSA. For each voxel in the AAL mask, we extracted the multivoxel activity patterns of 90 words within a sphere (radius = 8 mm) centered at that voxel. A neural 90 × 90 RDM was computed based on Pearson distance and was correlated with the semantic benchmark RDM while controlling for the two low-level RDMs of visual and phonological similarity of word stimuli, which produced a correlation coefficient for this voxel. By moving the searchlight center through the AAL mask, we obtained a correlation map for each participant. This map was Fisher-*z*-transformed and then spatially smoothed using a 6-mm FWHM Gaussian kernel. The correlation maps of native and delayed groups were compared using a two-sample *t*-test.

### Univariate semantic abstractness analysis in two deaf groups

#### GLM

The neural semantic abstractness effects were computed using the same GLMs in the “ROI definition” section. For each deaf participant, we computed the following three contrasts for ROI and whole-brain group comparisons using all the runs: abstract > concrete, abstract words > baseline, concrete words > baseline.

#### ROI analysis

For each ROI, we extracted the averaged beta values of abstract words > baseline and concrete words > baseline, from each deaf participant, and compared between two word types using one-tailed paired *t*-tests to test for the semantic abstractness effects in each deaf group. Two-way ANOVA was then adopted to examine group effects, with word type (concrete/object, abstract/nonobject) as a within-subject factor and group (native, delayed) as a between-subject factor.

#### fROI analysis

We also carried out individualized functional Region-of-Interest (fROI) analyses (Cohen et al., 2019; Murty et al., 2020), which examined semantic selectivity in individually defined functional voxels, with the fROI localization and selectivity calculation using independent datasets. Specifically, we estimated each deaf participant’s whole-brain beta and *t* maps for the contrast between abstract/nonobject and concrete/object words in odd and even runs. The semantic ROIs defined above in hearing participants were taken as anatomical constraints. In each hearing-group-ROI, for each deaf participant, we localized his/her top 50 selective voxels (i.e., voxels with highest *t* values to the contrast) in the odd runs, extracted the mean beta values of these voxels to abstract and concrete words, respectively, in the even runs. This procedure was repeated with fROI defined in the even runs and beta values calculated in the odd runs. The beta values were averaged across two iterations for each ROI in each participant and compared between two deaf groups using the abovementioned statistical analyses. We also repeated the fROI analyses at the fROI size of 100 voxels and obtained very similar results.

#### Whole-brain group comparison

The whole-brain semantic abstractness effects were examined in each deaf group, by one-sample *t*-tests on the whole-brain beta-weight images of abstract > concrete in each group. For the interaction between group (native, delayed) and word-type (abstract, concrete), the abstract > concrete beta-weight maps of two deaf groups were submitted to a two-sample *t*-test.

### Brain visualization

The brain results were projected onto the MNI brain surface for visualization using BrainNet Viewer (version 1.7; https://www.nitrc.org/projects/bnv/; (Xia et al., 2013)) with the default “interpolated” mapping algorithm, unless stated explicitly otherwise. Regional labels were derived based on the AAL template in xjview (by Xu Cui, http://www.alivelearn.net/xjview/).

## Data availability

Source data files have been provided for Table 1 and all the figures. Additional behavioral and neural data have been made available on OSF at the link https://osf.io/wz6q9/.

## Acknowledgments

We thank Dr. Xi Yu for helpful discussions on earlier drafts of the manuscript. We thank Ms. Yun Hao, Anran Deng, and Wei Liang for their assistance in data collection. This work was supported by the National Science and Technology Innovation 2030 Major Program (2021ZD0204104), the National Natural Science Foundation of China (31925020 and 82021004 to Y.B, 31700943 and 32171052 to X.W.), and Changjiang Scholar Professorship Award (T2016031 to Y.B.). The funders had no role in study design, data collection and analysis, decision to publish, or preparation of the manuscript.

## Supplementary files

- Supplementary file 1. Behavioral performances of two deaf groups in two reaction-time tasks.
- Supplementary file 2. Whole-brain RSA results of pixelwise similarity of visual words in two deaf groups.

---

**Figure 1-Figure supplement 1.**
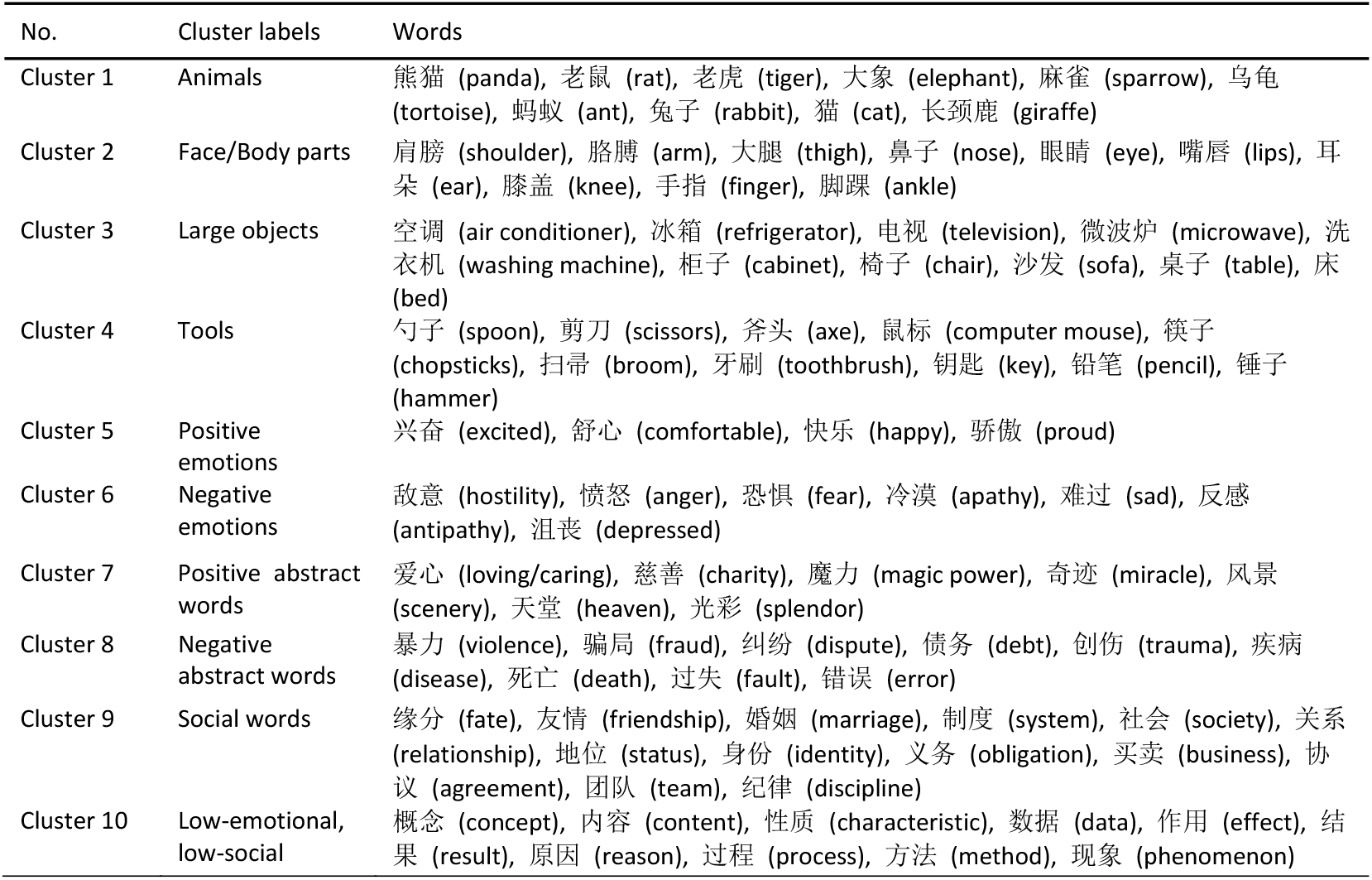
Ninety words in the fMRI task, grouped into ten semantic clusters based on k-means clustering of the group-mean hearing semantic space.

**Figure 1-Figure supplement 2.**
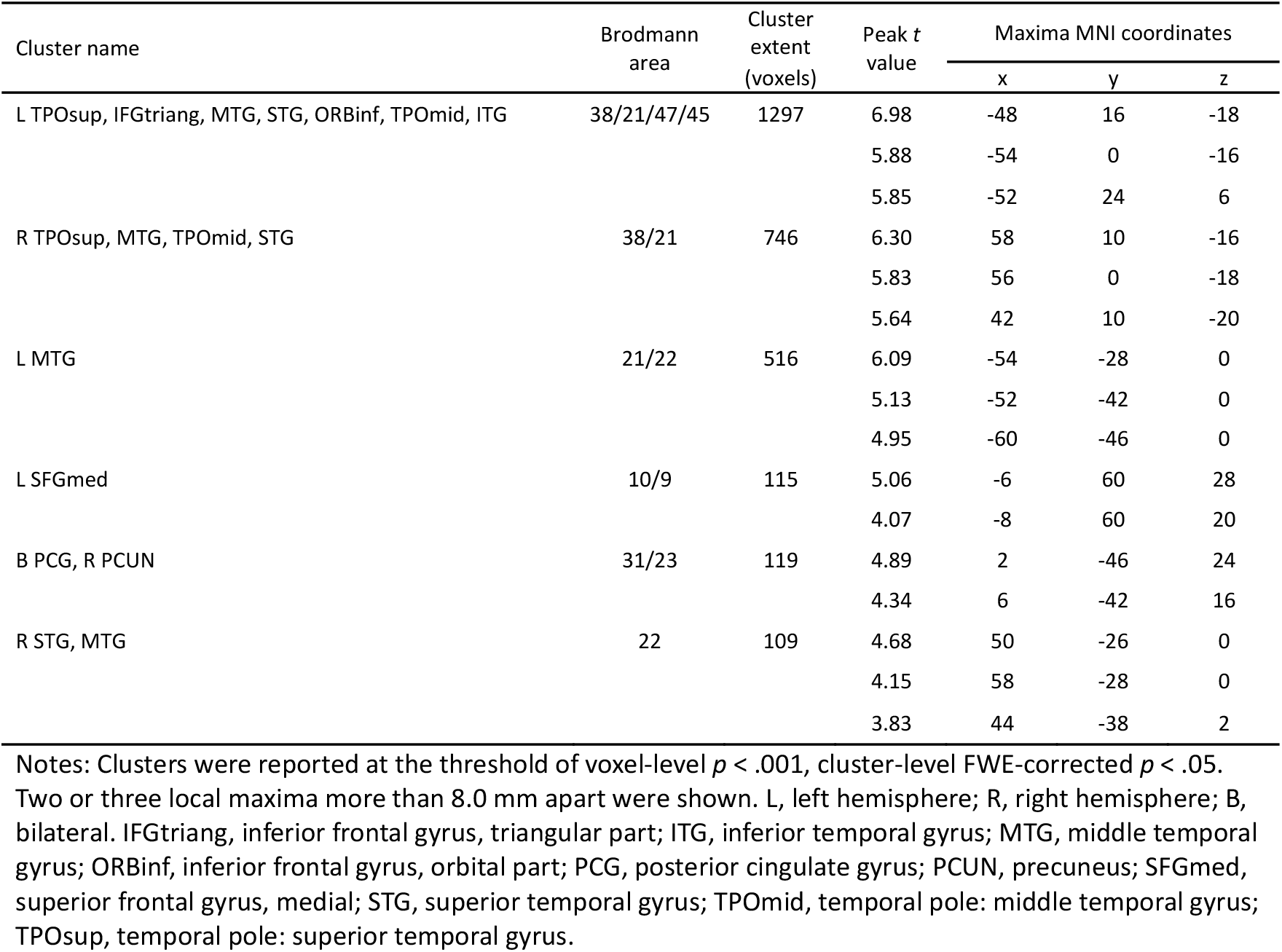
Whole-brain results of regions showing stronger activations to abstract/nonobject words than concrete/object words in 33 hearing participants.

**Figure 1-Figure supplement 3.**
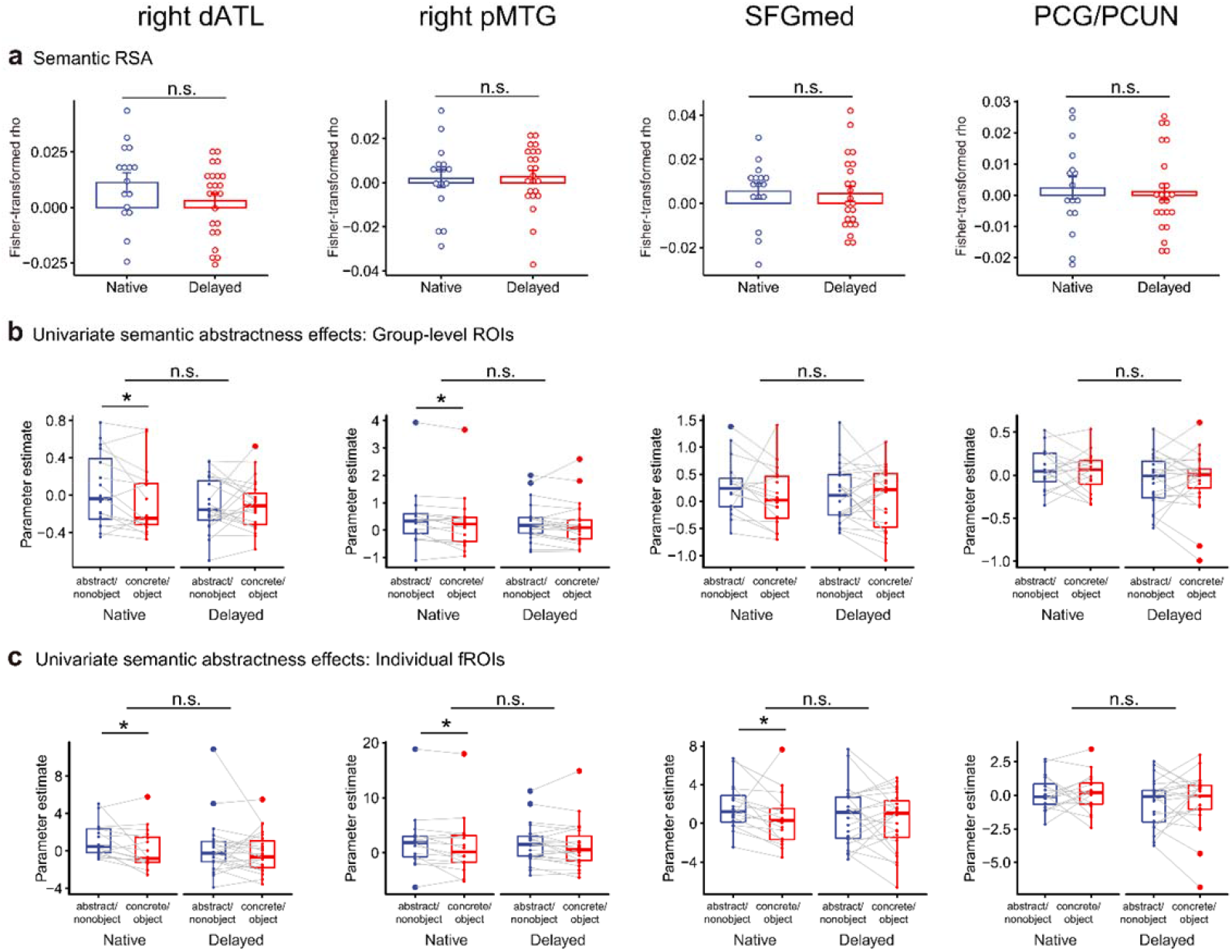
Additional ROI-level RSA (**a**) and univariate semantic abstractness results (**b** and **c**) in native and delayed signers. dATL, dorsal anterior temporal lobe; pMTG, posterior middle temporal gyrus; SFGmed, superior frontal gyrus, medial; PCG, posterior cingulate gyrus; PCUN, precuneus. n.s., not significant, *p* > .05; *, *p* < .05. n = 16 in the native group; n = 23 in the delayed group.

**Figure 2-Figure supplement 1.**
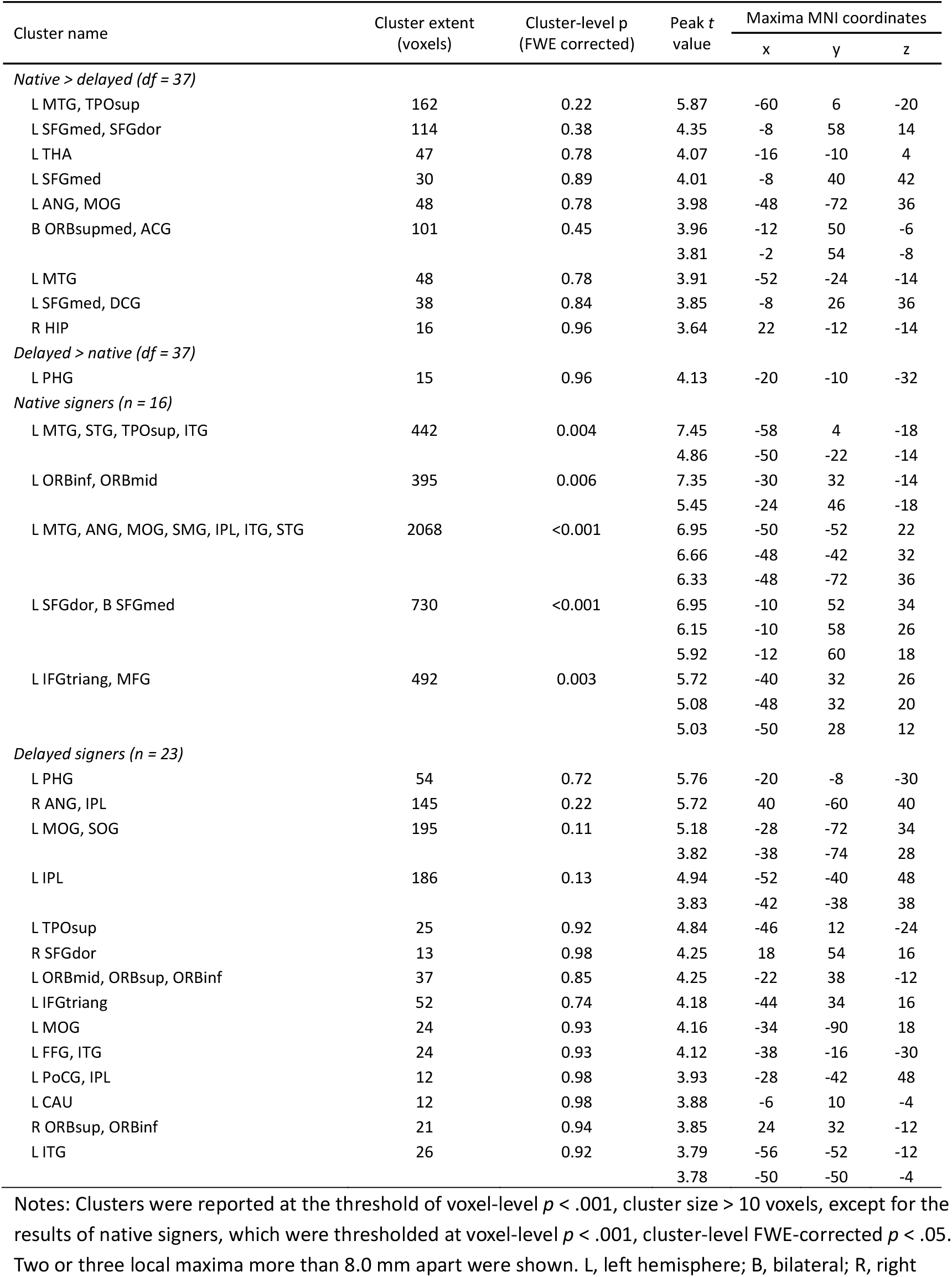
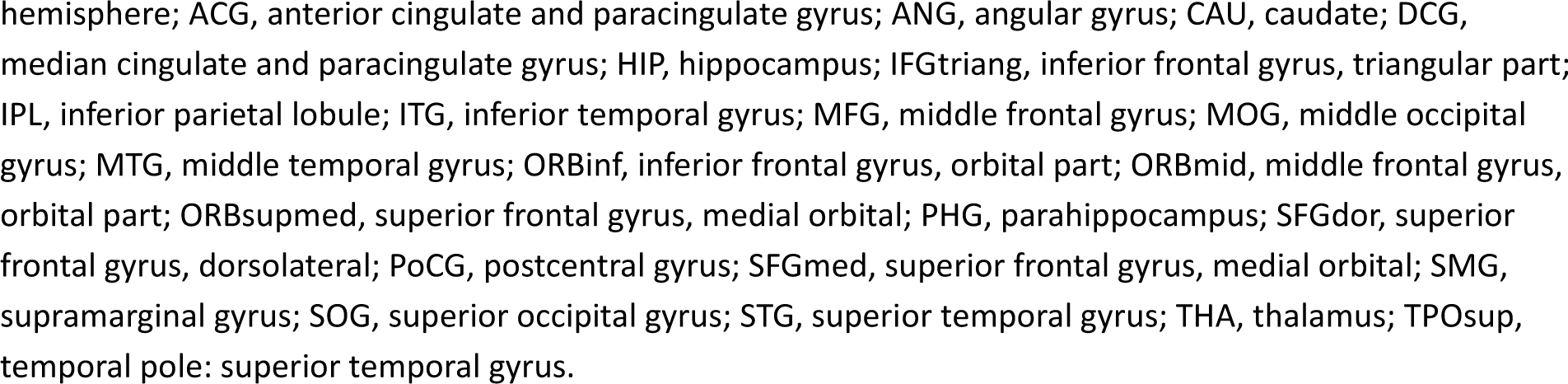
Whole-brain searchlight RSA results of semantic encoding.

**Figure 3-Figure supplement 1.**
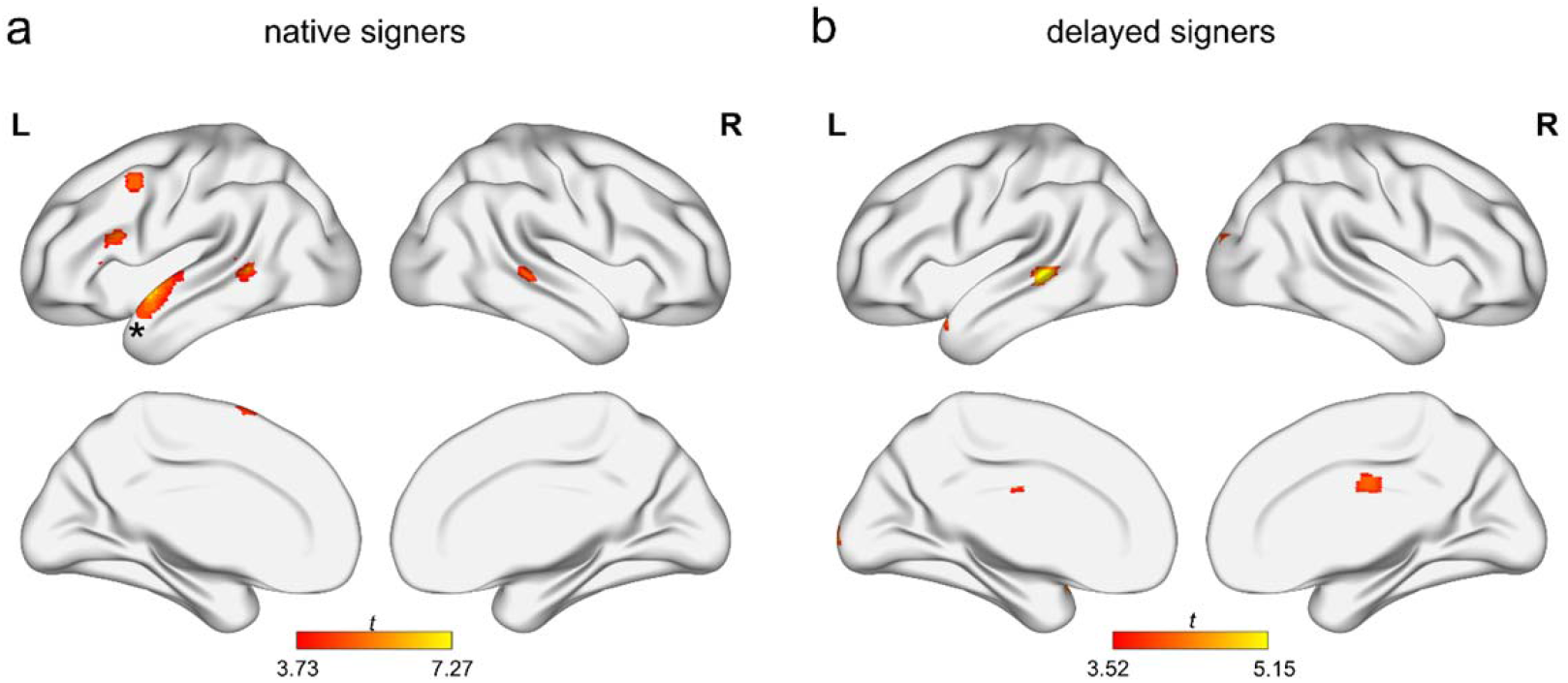
Whole-brain univariate contrast results of semantic abstractness in native (**a**) and delayed (**b**) signers. The neural effects were thresholded at voxel-level *p* < .001, cluster sizes > 10 voxels, and visualized using the “Maximum Voxel” mapping algorithm in BrainNet Viewer to illustrate small clusters. The asterisk in (a) indicates that the dATL cluster survived the cluster-level FWE-corrected *p* < .05. L, left hemisphere; R, right hemisphere. n = 16 in the native group; n = 23 in the delayed group.

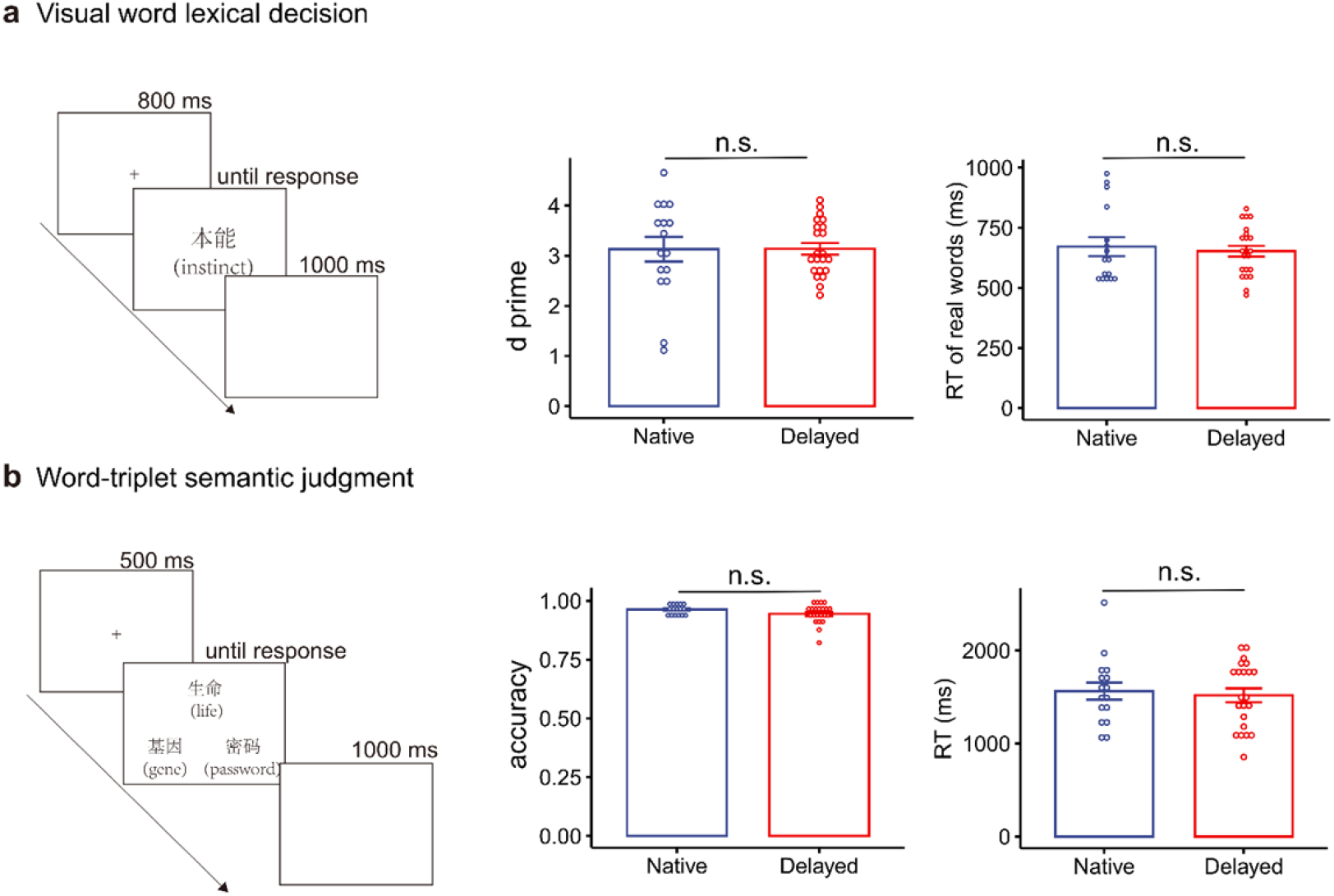

Native and delayed deaf signers did not significantly differ in written word processing in 2 behavioral reaction-time tasks. **a**. Lexical decision task, in which participants were asked to decide whether a 2-Chinese-character written stimulus denoted a real word or a pseudoword via button press. In each trial, a fixation point was presented for 800 ms, followed by a Chinese stimulus (font SONG, size 40). The stimulus disappeared upon the response and the next trial started 1 s later. The stimuli were 100 2-character Chinese nouns (word frequency, range: 0-1491 per million (Sun et al., 1997); number of strokes: range: 8-25) and 100 2-character pseudowords created by pseudo-randomly combining 2 characters of 100 real words. Bar plots show the d-prime and reaction time of real-word trials. d-prime was calculated using the Signal Detection Theory Calculator (version 1.2) provided by (Gaetano, 2017). **b**. Word-triplet semantic judgment task, in which participants were presented with three 2-character Chinese words and asked to decide which of the two choice words (bottom) was more semantically related to the probe word (top) via button press. In each trial, a fixation point was presented for 500 ms, followed by 3 Chinese words (font SONG, size 40), with the probe word presented at the top and two choice words arranged horizontally at the bottom. The stimuli disappeared upon the response and the next trial started 1 s later. The stimuli were 80 triplets of 2-character Chinese words (word frequency, range: 0 to 2340 per million; total number of strokes, range: 6-31), with all but four primarily used as nouns. Bar plots show overall accuracy and reaction time of correct trials. Both task procedures were implemented using E-prime 2 (Psychology Software Tools, Inc.; Pittsburgh, PA). n = 16 in the native group; n = 22 in the delayed group.

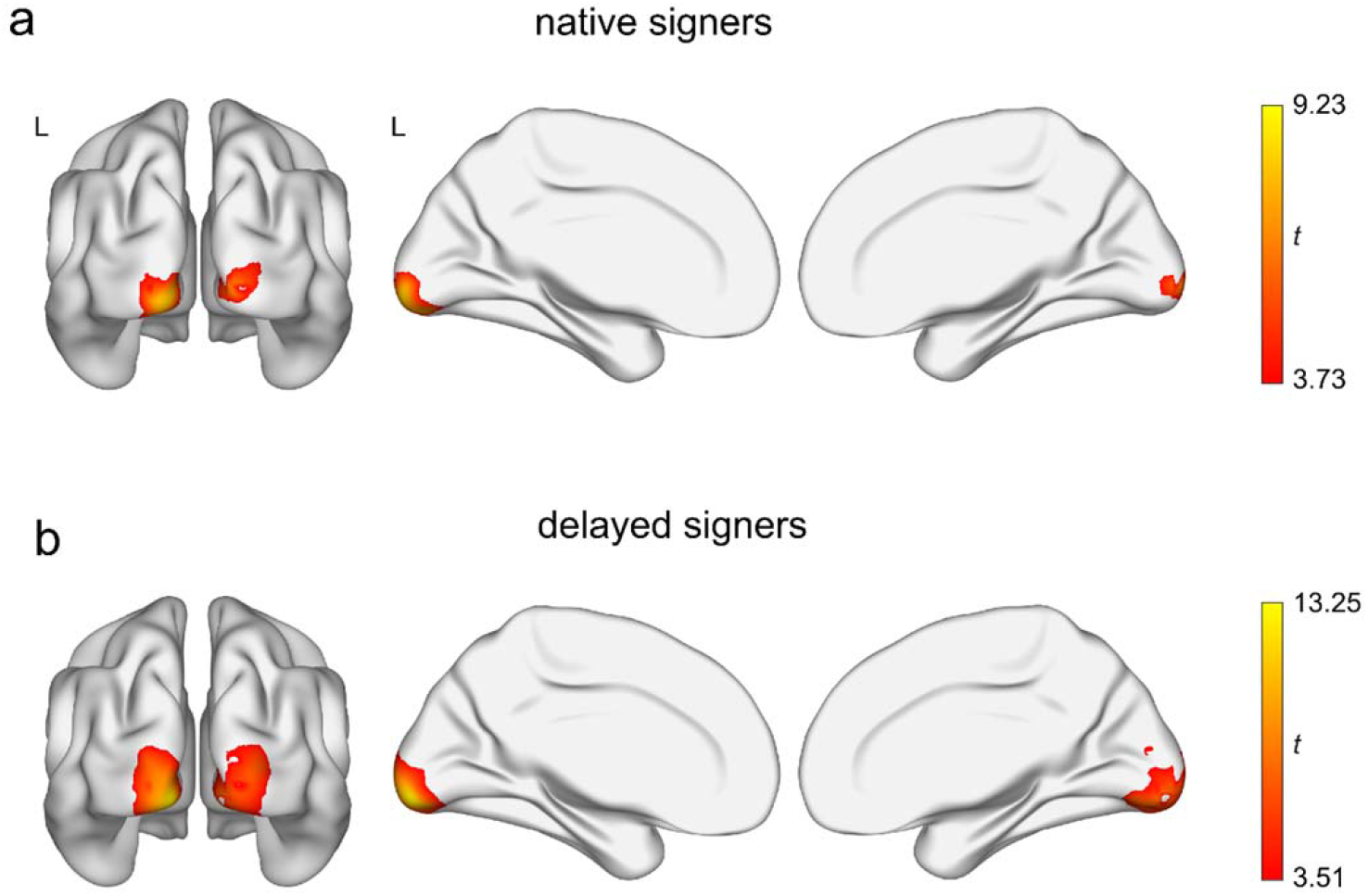

Whole-brain RSA results showed that pixelwise similarity of visual words were significantly encoded in early visual cortex in both native **(a)** and delayed **(b)** signers. The neural effects were thresholded at voxel-level *p* < .001, FWE-corrected *p* < .05. L, left hemisphere. n = 16 in the native group; n = 23 in the delayed group.

